# Cortical Responses to the Amplitude Envelopes of Sounds Change with Age

**DOI:** 10.1101/2020.10.23.352880

**Authors:** Vanessa C. Irsik, Ala Almanaseer, Ingrid S. Johnsrude, Björn Herrmann

## Abstract

Many older listeners have difficulty understanding speech in noise, when cues to speech-sound identity are less redundant. The amplitude envelope of speech fluctuates dramatically over time, and features such as the rate of amplitude change at onsets (attack) and offsets (decay) signal critical information about the identity of speech sounds. Aging is also thought to be accompanied by increases in cortical excitability, which may differentially alter sensitivity to envelope dynamics. Here, we recorded electroencephalography in younger and older human adults (of both sexes) to investigate how aging affects neural synchronization to 4-Hz amplitude-modulated noises with different envelope shapes *(ramped:* slow attack & sharp decay; *damped:* sharp attack & slow decay). We observed that subcortical responses did not differ between age groups, whereas older compared to younger adults exhibited larger cortical responses to sound onsets, consistent with an increase in auditory cortical excitability. Neural activity in older adults synchronized more strongly with rapid-onset, slow-offset (damped) envelopes, was less sinusoidal, and more peaked. Younger adults demonstrated the opposite pattern, showing stronger synchronization with slow-onset, rapid-offset (ramped) envelopes, as well as a more sinusoidal neural response shape. The current results suggest that age-related changes in the excitability of auditory cortex alter responses to envelope dynamics. This may be part of the reason why older adults experience difficulty understanding speech in noise.

**Significance Statement:** Many middle aged and older adults report difficulty understanding speech when there is background noise, which can trigger social withdrawal and negative psychosocial health outcomes. The difficulty may be related to age-related changes in how the brain processes temporal sound features. We tested younger and older people on their sensitivity to different envelope shapes, using EEG. Our results demonstrate that aging is associated with heightened sensitivity to sounds with a sharp attack and gradual decay, and sharper neural responses that deviate from the sinusoidal features of the stimulus, perhaps reflecting increased excitability in the aged auditory cortex. Altered responses to temporal sound features may be part of the reason why older adults often experience difficulty understanding speech in social situations.

## Introduction

Sensitivity to temporal features of sound, such as dynamic fluctuations in the amplitude envelope, is considered critical for speech intelligibility (Drullman et al., 1994; Shannon et al., 1995). Aging is associated with a decline in processing auditory temporal features (Gordon-Salant and Fitzgibbons, 1999) and speech intelligibility, particularly in the presence of background sounds (Gordon-Salant, 2006). In addition to peripheral hearing loss (presbycusis) (Frisina and Frisina, 1997) and cognitive decline (Wayne and Johnsrude, 2015; Griffiths et al., 2020), evidence increasingly suggests that poorer speech intelligibility in older individuals may be related to changes in how the cerebral cortex responds to amplitude envelopes (Millman et al., 2017; Goossens et al., 2018).

Neural activity readily synchronizes with lower-frequency (<20-Hz) sinusoidal amplitude envelopes of sounds (Aiken and Picton, 2008), but older adults often exhibit greater synchronization than younger to such amplitude modulations (AMs) (Goossens et al., 2016; Presacco et al., 2016a, 2016b; Herrmann et al., 2019). Enhanced AM synchronization may be disadvantageous for speech-in-noise perception (Millman et al., 2017; Goossens et al., 2018, 2019), because it may distort envelope pattern and depth cues (Moore and Glasberg, 1993; Schlittenlacher and Moore, 2016). Exaggerated AM synchronization may be related to heightened excitability of the auditory cortex in older people (Snyder and Alain, 2005; Bidelman et al., 2014; Herrmann et al., 2016; Salvi et al., 2017), perhaps resulting from reduced neural inhibition (Caspary et al., 2008).

The shape of amplitude envelopes in speech varies, including in the shape of the attack (rise) and decay (fall) portions (Rosen, 1992). Envelope-shape cues are important for identifying and discriminating between consonants (e.g., /pa/ versus /ta/) (van der Horst et al., 1999). Envelope shape can also alter cochlear excitation (Carlyon, 1996) and neural synchronization patterns (Pressnitzer et al., 2000; Lu et al., 2001; Neuert et al., 2001). Inferior colliculus neurons synchronize more strongly with damped (sharp attack, gradual decay) compared to ramped (gradual attack, sharp decay) envelope shapes in aged rats, whereas the opposite occurs for young rats (Herrmann et al., 2017). Synchronization to ramped and damped envelopes may also differ between older and younger human listeners: increased synchronization to sharp attacks in sounds may explain why older individuals report difficulty suppressing distracting sounds (Parmentier and Andrés, 2009; Mishra et al., 2014), and with speech-in-noise perception when modulated background sound (containing sharp attacks) is present (Moore and Glasberg, 1993; Millman et al., 2017).

Studies in humans and animals almost exclusively focus on synchronization at the stimulation frequency (Purcell et al., 2004; Dimitrijevic et al., 2016; Henry et al., 2017; Herrmann et al., 2017, 2018). Yet, energy is also commonly observed at the harmonics (Lins et al., 1995; Zhu et al., 2013), indicating responses are not fully sinusoidal (Dallos, 1973; Mayoral et al., 2017). Analysis of non-sinusoidal response features, like the harmonics, improves classification of neural synchronization in clinical settings (Cebulla et al., 2006) and predictions about AM coding using computational modeling (Vasilkov and Verhulst, 2019; Keshishzadeh et al., 2020). Further, non-sinusoidal signal-shape features – such as sharpness – can provide important information about neural dysfunction (Cole et al., 2017). Considering neural response features other than synchronization at the stimulation frequency may provide a better understanding of age-related neural synchronization changes.

In the present study, we examine neural synchronization to narrowband noise stimuli with ramped and damped envelope shapes in younger and older human adults. We use stimuli in two carrier-frequency bands (0.9-1.8 kHz, 1.8-3.6 kHz) which are within the frequency range to which human hearing is most sensitive, and in which the articulatory resonances that indicate speech sound identity (i.e., formants), are found. We also expand on traditional Fourier-based analyses to characterize non-sinusoidal response features.

## Materials and Methods

### Participants

Forty-nine younger (25 younger: 9 males and 16 females aged 18-32 years, M = 21.8 years, ± s.d. = 3.2 years) and older (24 older: 6 males and 18 females aged 50-83 years, M = 66.1 years, ± s.d. = 8.0) individuals were recruited for this experiment from the Western University Psychology subject pool and the surrounding community of London, Ontario (Canada) via Western’s neuroscience research registry (OurBrainsCAN; ourbrainscan.uwo.ca). All participants provided informed consent according to a protocol approved by Western’s Research Ethics Board (REB #112015), and either received course credit or financial compensation of $10 CAD per hour. The forty-nine participants included in this study reported having no hearing loss, hearing aid usage, neurological issues, or psychiatric disorders. Data from three additional individuals were not included due to a technical error during data recording (N=1), a neurological disorder (N=1), or hearing aid usage (N=1). Data from the younger participant group were also analyzed in the stimulus-selection phase of the experiment (Figure 2).

### Acoustic stimuli

Stimuli were narrowband noises generated by adding 150 randomly sampled carrier-frequency components with different onset phases from one of two possible carrier frequency bands (low: 0.9–1.8 kHz; high 1.8–3.6 kHz). Frequency bands were chosen to span the range of highest human sensitivity, and much of the energy that contributes to discrimination of speech sounds. Narrowband noises were amplitude modulated at a rate of 4 Hz with either a ramped (gradual attack and sharp decay) or damped (sharp attack and gradual decay) envelope shape (Figure 1; Herrmann et al., 2017). Amplitude envelopes were created by varying parameters of the following equation:

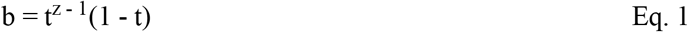

where *t* is a time vector representing one cycle (0.250 s), *z* determines the envelope shape, and *b* is the resulting function used to modulate the noise. A *z* parameter of 2 generates a symmetrical envelope shape, while a value closer to 1 generates an envelope with a damped shape (sharp attack and gradual decay). Varying the *z* parameter also impacts the sharpness and half-life of the oscillation. We used a *z* parameter of 1.4491 and 1.15 (based on Herrmann et al., 2017) to generate weakly and strongly modulated envelope shapes, respectively (Figure 1). Strongly modulated damped envelopes have sharp onsets and a 168.4 ms half-life, while weakly modulated envelopes have more sloped onsets and a 195 ms half-life. Weakly and strongly modulated ramped envelope shapes were created by mirroring the vector *b.* Stimuli were normalized relative to peak amplitude and presented at approximately 75 dB SPL (identical for all listeners).

**Figure 1.**
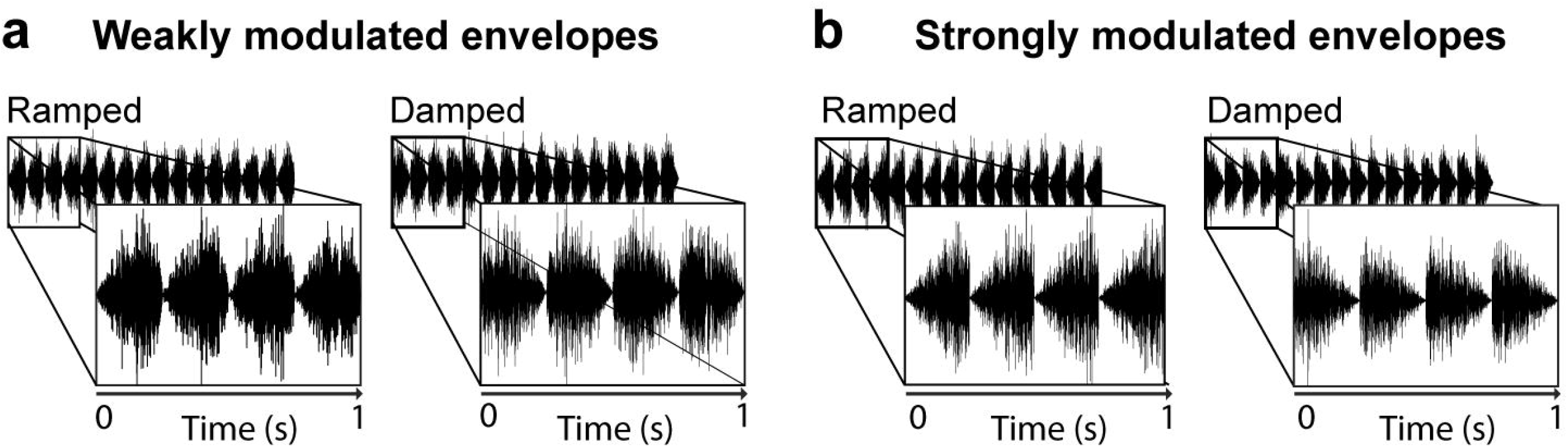
Samples of acoustic stimulation with different envelope shapes. Ramped shape: gradual attack and sharp decay; and damped shape: sharp attack and gradual decay. Narrowband noises were amplitude modulated at a rate of 4 Hz with either **(a)** weakly ramped/damped or **(b)** strongly ramped/damped envelopes.

In an effort to select either weakly or strongly modulated envelope shapes for the main experiment which investigates response changes with age, we examined neural synchronization to either weakly or strongly ramped and damped 4-Hz amplitude-modulated noise stimuli in two groups of younger adults. One group of participants listened to noises with weakly ramped/damped envelope shapes, while the other group listened to noises with strongly ramped/damped envelope shapes. Additional details regarding the participants, procedure, and analyses are reported in the caption of Figure 2. We observed that synchronized neural activity was larger for strong compared to weak envelope shapes. Therefore, we utilize strongly shaped envelopes to investigate whether neural synchronization with ramped and damped amplitude modulations changes with age.

**Figure 2.**
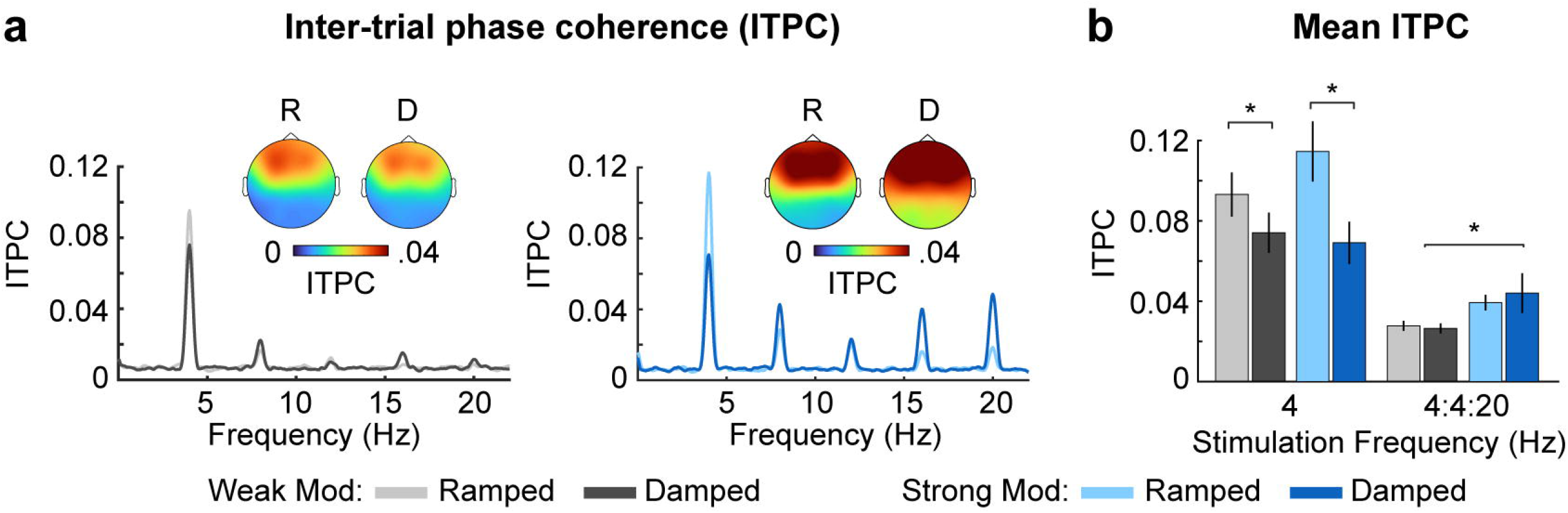
Stimulus selection experiment. **Method:** Data from two groups of younger subjects were analyzed as part of a stimulus selection experiment. One group listened to narrowband noises with weakly ramped/damped envelope shapes (N=25; 18 females; age range: 18-25 years, M = 20.2 years, ± s.d. = 2.3 years), while the other group listened to noises with strongly ramped/damped envelope shapes (N=25; 16 females; age range: 18-32 years, M = 21.8 years, ± s.d. = 3.2 years) and EEG was recorded as described in the Materials & Methods section. **Analysis:** Neural synchronization was analyzed using identical methods as the inter-trial phase coherence (ITPC) analysis of the main experiment. To examine whether neural synchronization strength differed as a function of envelope shape strength, ITPC at the amplitudemodulation frequency (4 Hz) and for the 4-Hz fundamental/harmonic series (4:4:20 Hz) were submitted to separate ANOVAs, each with envelope shape (ramped, damped) and carrier frequency (low, high) as within-subject factors and envelope shape strength (weak, strong) as a between-subjects factor. **Results:** The results indicate that synchronized neural activity was larger for strong compared to weak envelope shapes (effect of shape strength: *F*_1,48_ = 4.44, *p* = .04, η^2^_p_ = .09), but this effect was only observed when considering responses to the harmonic series (4:4:20 Hz). **Figure: (a)** Inter-trial phase coherence (ITPC), and **(b)** mean ITPC at the stimulation frequency (4 Hz) and at the 4-Hz fundamental/harmonic series (4:4:20 Hz) are plotted as a function of envelope shape strength (weak, strong) and envelope shape (ramped, damped). Topographies in **(a)** reflect mean ITPC at the 4-Hz stimulation frequency and are shown for each envelope shape (ramped, damped) and age group (younger, older). Error bars reflect standard error. *p < 0.05.

### Task procedure

The experiment was conducted in a single-walled sound-attenuating booth (Eckel Industries). Sounds were delivered through Sennheiser (HD 25 Light) headphones, using an RME Fireface 400 external soundcard controlled by a PC (Windows 10) and Psychtoolbox (Version 3) in MATLAB (R2017b). EEG was recorded while participants passively listened to a series of strongly modulated ramped and damped sounds while watching a muted captioned movie of their choice. Each stimulus had a duration of 4 s and stimuli were presented at an onset-to-onset interval of 5.021 s. Participants heard 28 ramped and 28 damped stimuli in each of the two carrier-frequency bands (low: 0.9–1.8 kHz; high: 1.8–3.6 kHz) during each of the 6 blocks, for a total of 168 trials per condition per person.

### Experimental design and statistical analysis

Statistics were conducted using a combination of IBM SPSS Statistics for Windows (v24) and MATLAB. Details of the specific variables and statistical tests for each analysis can be found in subsequent analysis subsections. In general, group differences were examined either using an analysis of variance (ANOVA) or independent-samples *t*-tests. Non-parametric Mann-Whitney U tests were used to analyze behavioral ratings, as these data were ordinal, not continuous. Significant main effects and interactions were followed up using */*-tests, with multiple comparisons corrected using the false discovery rate (FDR; Benjamini and Hochberg, 2016) correction. FDR corrected p-values are referred to as *P_FDR_*. Effect sizes are reported as partial eta squared (η^2^_p_) for ANOVAs and r_equivalent_ (r_e_; Rosenthal and Rubin, 2003), for *t*-tests. This experiment was not preregistered. Data are available at the project website on the Open Science Framework (OSF; https://osf.io/eq45x/).

### Behavioral hearing assessment

Pure-tone thresholds were measured for all participants at octave frequencies between 0.25 and 8 kHz in the left and right ear (see Figure 3a). Pure-tone thresholds were used to calculated pure-tone averages (PTA) across octave frequencies from 0.5 to 4 kHz (averaged across ears), to characterize the presence of hearing loss in a range of frequencies relevant to the stimuli from the main portion of the experiment (see General Methods and Materials). Average PTA thresholds were submitted to an independent-samples *t*-test with age group (younger, older) as the grouping variable.

**Figure 3.**
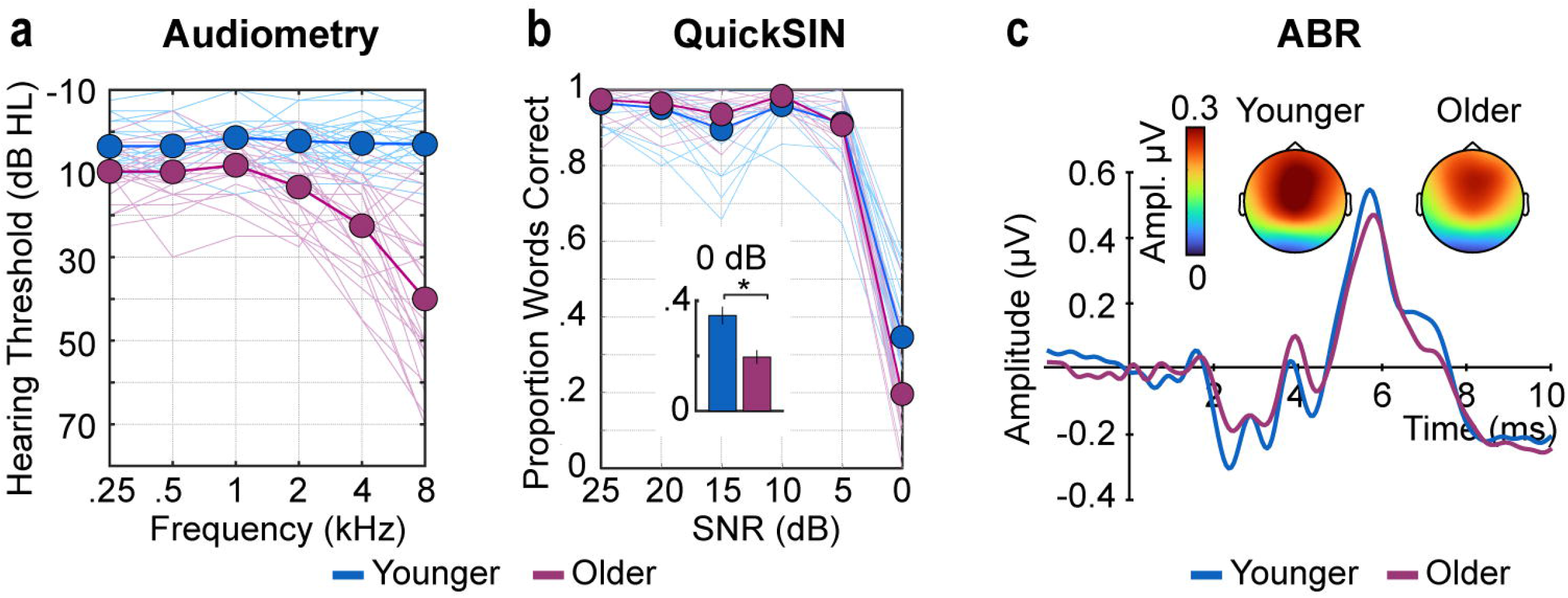
Hearing assessment measures. **(a)** Audiometric thresholds **(b)** QuickSIN performance, and **(c)** auditory brainstem responses (ABR) are plotted for each age group (younger, older). Topographies reflect the mean Wave V response. Thin lines in **(a)** and **(b)** reflect individual participants. Thick lines reflect the average across participants. Error bars reflect standard error. *p < 0.05.

Participants also answered questions taken from the Speech, Spatial, and Qualities of Hearing Scale (Gatehouse & Noble, 2004), asking them to use a Likert scale (0: ‘not at all’ to 10: ‘perfectly’) to rate their ability in listening situations requiring spatial hearing (N=2), speech perception in noise (N=2), and suppression of distracting background sounds (N=1). Average scores were generated for listening-situation categories with multiple questions (i.e., spatial hearing, speech perception in noise). Given that SSQ scores are ordinal, not continuous, we used separate Mann-Whitney U tests (non-parametric) examine age group (younger, older) differences on each listening situation category (spatial hearing, speech perception in noise, distractor suppression).

Participants completed the Quick Speech in Noise test (QuickSIN) (Killion et al., 2004), a clinical measure used to assess speech understanding in noise. All target sentences and babble noise were taken from the QuickSIN database. During the test, a target sentence, spoken by a female talker, was presented with four-talker babble as background noise (overall 70 dB SPL). Participants were instructed to listen to each sentence and type the words that they heard. Sentences were presented in sets of 6, which began with a 25-dB signal to noise ratio (SNR) and reduced in 5 dB steps until the final sentence was completed. Participants were each asked to complete 4 sentence sets (24 total sentences) that were randomly selected from 12 possible sets. We calculated performance for each SNR separately (25, 20, 15, 10, 5, 0 dB), and report the total proportion of correct words for each SNR. Performance was at ceiling for all SNRs except 0 dB. We therefore examined age group (younger, older) differences on performance at the 0 dB SNR using an independent-samples *t*-test.

### EEG recording and preprocessing

EEG was recorded from 16 active electrodes (Ag/AgCl) placed on the scalp using an electrode cap with spacing according to the 10/20 system (Biosemi ActiveTwo system). We also recorded and averaged signals from both mastoids to re-reference the data during offline analysis. During data recording, all electrodes were referenced to a feedback loop formed of two electrodes, a common mode sense (CMS) active electrode and a driven passive electrode (see www.biosemi.com/faq/cms&drl.htm). EEG was recorded at 16,384 Hz to target peripheral and subcortical sources during ABR recording (online low-pass filter 3334 Hz) and at 1024 Hz to isolate primarily cortical sources during envelope tracking (online low-pass filter of 208 Hz). All pre-processing was carried out offline using MATLAB software and the Fieldtrip toolbox (Oostenveld et al., 2011).

For ABR recordings, data were re-referenced to the averaged signal from both mastoids, a notch filter was used to attenuate signal at line-noise frequencies (60 Hz and 120 Hz), and then the EEG data were high-pass (80 Hz, 2743 points, Hann window) and low-pass filtered (2000 Hz, 101 points, Hann window). Continuous data were segmented into 12-ms epochs ranging from −2 ms to 10 ms time-locked to click onset. Epochs in which signal changed by more than 25 μV during the 0–10 ms time window in any channel were rejected (average rejection rate: 16 %).

For cortical EEG, data were re-referenced to the averaged signal from both mastoids and then high-pass (0.7 Hz, 2449 points, Hann window) and low-pass filtered (30 Hz, 101 points, Hann window). The continuous EEG data were segmented into epochs ranging from −0.5 to 4 s, time-locked to the onset of each stimulus. Ocular artifacts were removed using independent components analysis (Makeig et al., 1996). Epochs in which the signal changed by more than 150 μV in any channel were rejected (average rejection rate: 5%).

### EEG analysis: Peripheral and subcortical neural responses

We recorded click-evoked auditory brainstem responses (ABR) to derive an objective physiological measure of auditory peripheral and subcortical function. Participants were asked to passively listen to a series of isochronous clicks presented monaurally to the right ear, while watching a muted captioned movie of their choice and electroencephalography (EEG) was recorded. Each click had a 0.1 ms duration (rectangular window) and was presented monaurally to the right ear with an 11.3-ms onset-to-onset interval and an approximate sound level of 88 dB SPL. A total of 4000 clicks were presented with click polarity inverted on half of the trials, resulting in an equal proportion of condensation and rarefaction clicks.

A small subset of electrodes were used for the analysis to approximate a vertical electrode montage (Cz referenced to mastoid ipsilateral to sound presentation); this subset was chosen because it is known to maximize appearance of both Wave 1 and Wave V (Picton, 2010a). Peak latency was identified as the time point corresponding to maximum amplitude within a time window specific to Wave 1 (1–3 ms) and Wave V (5–7 ms). Peak amplitude was calculated by averaging the amplitude within a 0.5 ms window centered on Wave 1 or Wave V latency. Resulting peaks were visually inspected to ensure the response peaks well-characterized Wave 1 and Wave V responses.

No discernible Wave I or V peak could be identified for two individuals in the younger age group, both of whom required that more than 75% trials be rejected due to excessive artifact. These individuals were excluded from ABR analysis. Wave I and V amplitudes and latencies were calculated and then analyzed for the remaining participants (24 older and 23 younger) using 4 separate independent-samples t-tests, each which had age group (younger, older) as the grouping variable.

### EEG analysis: Cortical responses to sound onset

Single-trial time courses for each envelope shape (ramped, damped) and carrier-frequency band (low, high) were averaged separately. We examined P1 and N1, both of which are sensory-evoked responses with primary sources originating in auditory cortex (Hari et al., 1982; Näätänen and Picton, 1987; Pantev et al., 1988; Liégeois-Chauvel et al., 1994; Yoshiura et al., 1995), by averaging responses across a fronto-central electrode cluster that is sensitive to both responses (Fz, F3, F4, Cz, C3, C4) (Näätänen and Picton, 1987; Picton, 2010b). Mean amplitude was calculated by finding the time point corresponding to either the maximum (P1) or minimum amplitude (N1 peak) within a time window specific to the onset response (P1: 0.045-0.065 s; N1: 0.085-0.115 s), and then averaging the amplitude values within a corresponding averaging window (P1: 0.02-s; N1: 0.03-s) centered on the response peak. Visual inspection was performed to ensure the response peaks were accurately found for the P1 and the N1, for both ramped and damped envelope shapes.

We used onset response amplitude (P1, N1) as a metric of neural responsiveness to sound (cf. Snyder and Alain, 2005; Alain et al., 2014; Herrmann et al., 2016, 2019; Henry et al., 2017). P1 and N1 amplitudes were submitted to separate ANOVAs with envelope shape (ramped, damped) and carrier frequency (low, high) as within-subject factors and age group (younger, older) as a between-subjects factor.

### EEG analysis: Time-course correlation similarity

In order to better understand whether, and to what extent, cortical time courses differ between age groups, we quantified the degree of similarity between the neural time courses. After excluding the first 0.5 s (onset response range), we averaged responses across electrodes (Fz, F3, F4, Cz, C3, C4) and the two carrier-frequency band conditions (because no interactions with age group were observed when including carrier frequency as a factor, *F* < .44, *p* > .51, η^2^_p_ < .009), resulting in one averaged time course for each envelope shape (ramped, damped) condition and participant. Correlations between the averaged time courses were calculated, separately for ramped and damped stimuli, such that each participant’s time course was correlated with the time course of each participant within their ‘own’ age group and with the time course of each participant from the ‘other’ age group. For each envelope shape and each participant, the set of r values resulting from the correlations with time courses from other participants were categorized (‘own’ vs ‘other’ group) and averaged separately, yielding four mean correlations for each participant: two for each envelope shape: one for ‘own group’ and one for ‘other’ group. Larger own-group *r* values would indicate an individual’s response time course was highly synchronous with others in their own age group, while larger other-group *r* values would indicate an individual’s response time course was more synchronous with individuals in the other age group.

To quantify the degree of similarity between the neural time courses for younger and older adults, we compared average *r* values using an ANOVA with envelope shape (ramped, damped) and correlation type (own-group *r*, other-group *r*) as within-subjects factors and age group (younger, older) as the between-subjects factor.

### EEG analysis: Neural synchronization strength

In order to characterize neural synchronization to amplitude modulation in the narrowband noise stimuli, we calculated inter-trial phase coherence (ITPC) (Lachaux et al., 1999). For each condition, a fast Fourier transform was calculated for the 0.5 to 4 s time window of single-trial time courses (Hann window; zeropadding). The first 0.5 s were excluded from the analysis to prevent onset responses from affecting the neural-synchronization analysis. Each complex number resulting from the fast Fourier transform was divided by its absolute value and averaged across trials. ITPC values were derived by calculating the absolute value of the resulting average. ITPC can take on values between 0 and 1, with larger values indicating greater coherence. For each condition, ITPC was averaged across electrodes (Fz, F3, F4, Cz, C3, C4). Average ITPC was extracted at the amplitude modulation frequency (4 Hz; averaging window: ±0.05 Hz).

The fast Fourier transform decomposes a time-domain signal into unique sinusoidal components (co-sines), regardless of whether the time-domain signal has a dominant sinusoidal structure. As a result, any complex signals that contain periodic, but non-sinusoidal structure in the time-domain will yield peaks in the spectrum at the fundamental frequency as well as at the harmonics (Mayoral et al., 2017). Ignoring the response magnitude at the harmonics may leave out important information, since neural synchronization to amplitude-modulated sounds is often non-sinusoidal (Dallos, 1973; Lins et al., 1995; Cebulla et al., 2006; Zhu et al., 2013). Given that responses are clearly visible in the spectrum at the harmonics of the stimulation frequency (see Figures 2 and 4) we also averaged ITPC values at the stimulation frequency (4 Hz) and harmonics up to 20 Hz (i.e., 4, 8, 12, 16 and 20 Hz; abbreviated hereafter using 4:4:20 Hz; averaging window: ±0.05 Hz). By doing so, we can explore whether including non-sinusoidal response features, such as responses to the harmonics, offers additional information above and beyond what is observed at ITPC to the stimulation frequency (4 Hz).

To examine whether neural synchronization strength differed between age groups, ITPC at the amplitude-modulation frequency (4 Hz) and for the 4-Hz fundamental/harmonic series (4:4:20 Hz) were submitted to separate ANOVAs, each with envelope shape (ramped, damped) and carrier frequency (low, high) as within-subject factors and age group (younger, older) as a between-subjects factor.

### EEG analysis: Quantification of non-sinusoidal response patterns and signal shape

A growing body of evidence suggests that non-sinusoidal activity features can provide important information about underlying neural response properties (Cole and Voytek, 2017; Cole et al., 2017) including those that may indicate neural dysfunction (Sherman et al., 2016). We therefore investigated the degree to which synchronized neural responses diverge from a sinusoidal shape in two unique, but complementary, ways.

First, we investigated the harmonic structure of the ITPC frequency spectrum. High amplitude at the harmonics of the fundamental frequency would indicate that responses are less sinusoidal (Dallos, 1973; Mayoral et al., 2017). To approximate this, we extracted ITPC at the 4 Hz fundamental frequency (4 Hz, averaging window: ±0.05 Hz), and at the frequency of the harmonics from 8 to 20 Hz (i.e., 8, 12, 16 and 20 Hz; 8:4:20 Hz, averaging window: ±0.05 Hz) for each condition, and calculated the ratio between the two according to the following equation:

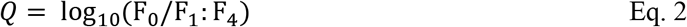

where F_0_ refers to mean ITPC at 4 Hz, and F_1_:F_4_ refers to mean ITPC across harmonic frequencies: 8, 12, 16 and 20 Hz. Larger *Q* values indicate a more sinusoidal synchronization response, whereas smaller *Q* values indicate a more non-sinusoidal synchronization response. *Q* was submitted to an ANOVA with envelope shape (ramped, damped) and carrier frequency (0.9–1.8 kHz, 1.8–3.6 kHz) as within-subjects factors and age group (younger, older) as the between-subjects factor.

Using a second approach, we quantified specific non-sinusoidal signal shape features. For this analysis, the amplitude values of trial-averaged time courses (averaged across electrodes: Fz, F3, F4, Cz, C3, C4) were related to the 4-Hz stimulus phase. That is, the time-course amplitude data were binned according to phase values assuming a 4-Hz sinusoid (number of bins: 100; window width: 0.063 radians), such that signal amplitude was represented as a function of phase (Figure 7a). An exponential cosine function was fit to the amplitude data using the following equation:

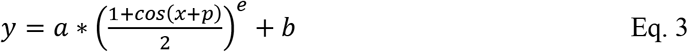

where *y* is the vector of binned amplitudes as a function of phase, *a* is the parameter for amplitude, *x* is the starting phase value, *p* is a vector of the 100 linearly spaced phase values relating to amplitude values in *y, b* is the intercept, and *e* is the exponent parameter which determines the sharpness of the function (see Figure 7b). An exponent of 1 reflects a sinusoid. An exponent larger than 1 means the signal is non-sinusoidal, and the function increases in sharpness with increasing exponent. Here, we analyzed two parameters from each fit, amplitude and exponent (sharpness), to directly quantify whether neural responses are hyper-responsive (amplitude) and contain non-sinusoidal response features (exponent) (note that *b* – the intercept – is not meaningful here as it is close to zero due to the high-pass filter). The estimated amplitude *a* and the estimated exponent *e* were submitted to an ANOVA with envelope shape (ramped, damped) and carrier frequency (low, high) as within-subjects factors and age group (younger, older) as the between-subjects factor. The absolute value of the fitted amplitude *a* was calculated prior to the ANOVA, because the inclusion of *e* in the formula sometimes led to a sign inversion of *a*.

## Results

### Younger and older listeners differ in behavioral hearing assessment, but not in subcortical responses

Pure-tone thresholds for octave frequencies between 0.25 and 8 kHz (averaged across ears) are plotted in Figure 3a. All participants had pure-tone average (PTA) thresholds (0.5 to 4 kHz averaged across ears) less than or equal to 31 dB HL. Relative to younger participants, older adults had elevated PTA thresholds (+10.93 dB HL; t_47_ = −6.96, *p* = 9.56 × 10^-9^, r_e_ = 0.71) and lower self-reported ratings for spatial hearing (U = 183.5, *p* = .018), sound distractor suppression (U = 154, *p* = .003), and understanding speech in the presence of background noise (U = 154.5, *p* = .003) (Gatehouse and Noble, 2004).

Older and younger adults performed at ceiling on the QuickSIN except for the most difficult SNR level (0 dB). At 0 dB, both groups exhibited proportions of correctly reported words that were significantly lower than 1 (younger: M = 0.34, s.e. = 0.03, 95% CI [0.28 0.40]; older: M = 0.20, s.e. = 0.02, 95% CI [0.15 0.24], and proportions were lower for older compared to younger adults (0 dB, t_47_ = 3.66, *p* = .001, r_e_ = 0.47; Figure 3b.

Despite these age-group differences in behavioral assessment metrics, at the neural level, there was no group difference in Wave I or Wave V amplitude (Wave I: *p* = .823; Wave V: *p* = .295) or latency (Wave I: *p* = .105; Wave V: *p* = .574) in response to click stimulation (11.3 Hz, 88 dB SPL) (Figure 3c). To examine whether hearing loss, instead of age, was associated with a reduction in subcortical function, we calculated the partial correlation (controlling for age) between average PTA thresholds and the ratio between Wave V and I amplitude (Wave V/I ratio), but did not find a significant relationship (*r*_44_ = −.1, *p* = .51). These findings do not suggest that auditory nerve and subcortical function differed across age groups.

### Aged auditory cortex is hyper-responsive to sound

Aging and hearing loss are associated with maladaptive cortical plasticity that leads to greater responsivity to stimulation by neural populations in auditory cortex (Snyder and Alain, 2005; Alain et al., 2014; Auerbach et al., 2014; Herrmann et al., 2016, 2017, 2018; Henry et al., 2017; Salvi et al., 2017; Herrmann and Butler, 2020). Hyper-responsiveness may be an index of reduced inhibition in cortical circuits (Caspary et al., 2008; Knipper et al., 2013; Ng and Recanzone, 2018). In order to test whether the cortex in the sample of older adults tested here is hyper-responsive, we compared neural responses elicited by sound onset between age groups (Figure 4a).

**Figure 4.**
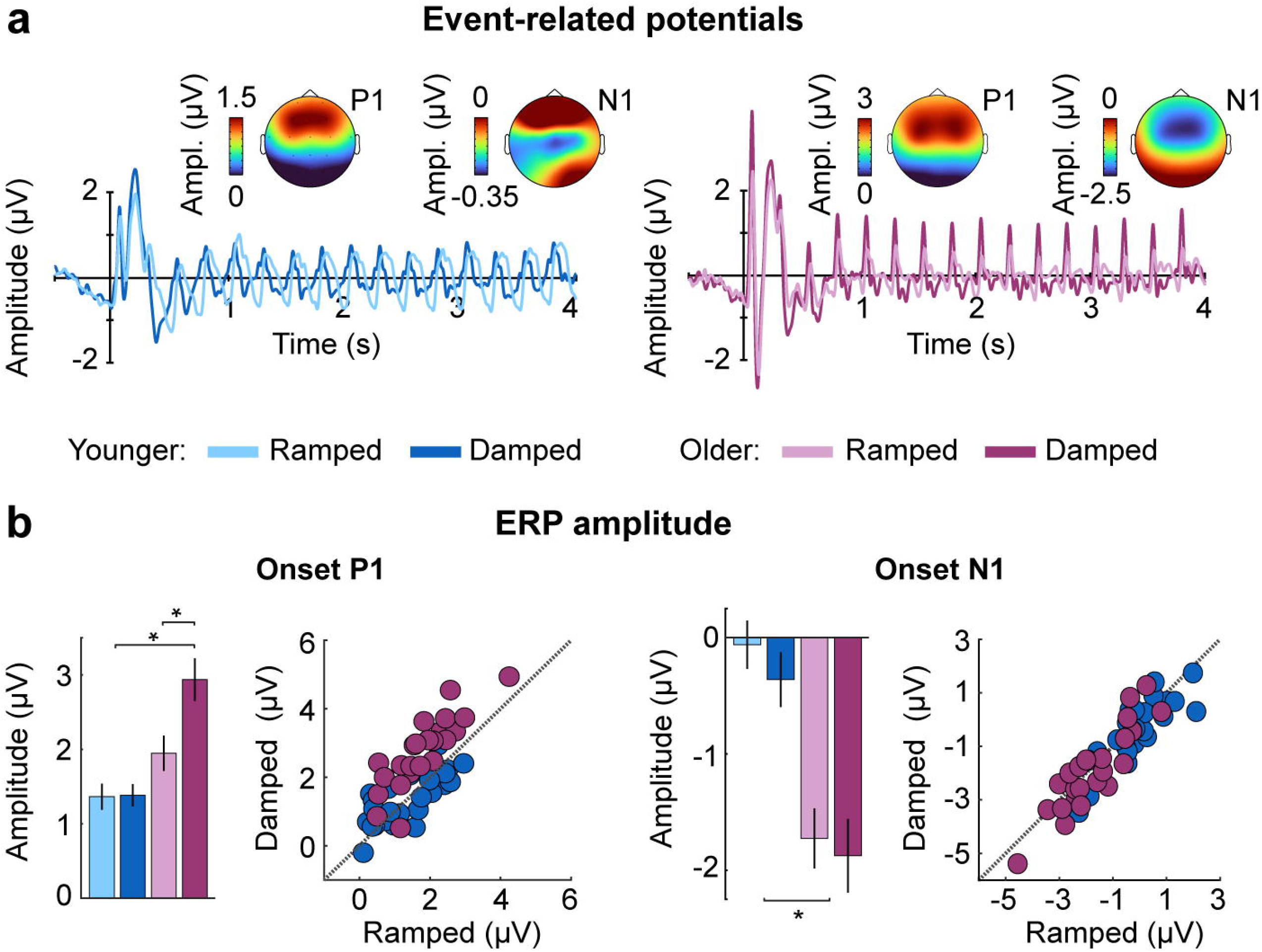
Neural responses to amplitude-modulated noises with different envelope shapes for younger and older adults. **(a)** Neural time courses, **(b)** P1 amplitude (left panel), and N1 amplitude (right panel) are plotted for each age group (younger, older) and envelope shape (ramped, damped). Topographies in **(a)** reflect mean P1 or N1 amplitude (averaged across envelope shapes and carrier frequency bands) for each age group (younger, older). Error bars reflect standard error. *p < 0.05.

Response amplitudes for both P1 and N1 were larger for older compared to younger adults (effect of age group: [P1: *F*_1,47_ = 13.18, *p* = .001, η^2^_p_ = .22] [N1: *F*_1,47_ = 20.68, *p* = 3.8 × 10^-5^, η^2^_p_ = .31]; see Figure 4b), indicating hyper-responsiveness to sound. Stimuli with damped envelope shapes also elicited larger P1 and N1 amplitudes compared to noises with ramped envelope shapes (effect of envelope shape: [P1: *F*_1,47_ = 35.73, *p* = 2.9 × 10^-7^, η^2^_p_ = .43] [N1: *F*_1,47_ = 5.36, *p* = .025, η^2^_p_ = .10]). This is consistent with previous research showing that N1 amplitude is larger when the stimulus rise time is fast (Picton, 2008), probably because sharp onsets drive more synchronous activity than slower onsets. Finally, there was an envelope shape × age group interaction for P1 (*F*_1,47_ = 32.89, *p* = 6.8 9 × 10^-7^, η^2^_p_ = .41). Older adults showed larger P1 amplitudes for damped compared to ramped envelopes (t_23_ = 8.22, *P_FDR_* = 5.3 × 10^-8^ .013, r_e_ = .86), while younger adults showed no such difference *(p_FDR_* = .864). None of the other effects or interactions were significant (*F* < 3.8,*p* > .05, η^2^_p_ < .1).

### Neural time courses differ between younger and older adults

In order to quantify differences in the response time courses between older and younger adults, we calculated correlations between individual participants’ response time courses within and across age groups, separately for stimuli with ramped (Figure 5a top panel) and damped envelope shapes (Figure 5a bottom panel).

**Figure 5.**
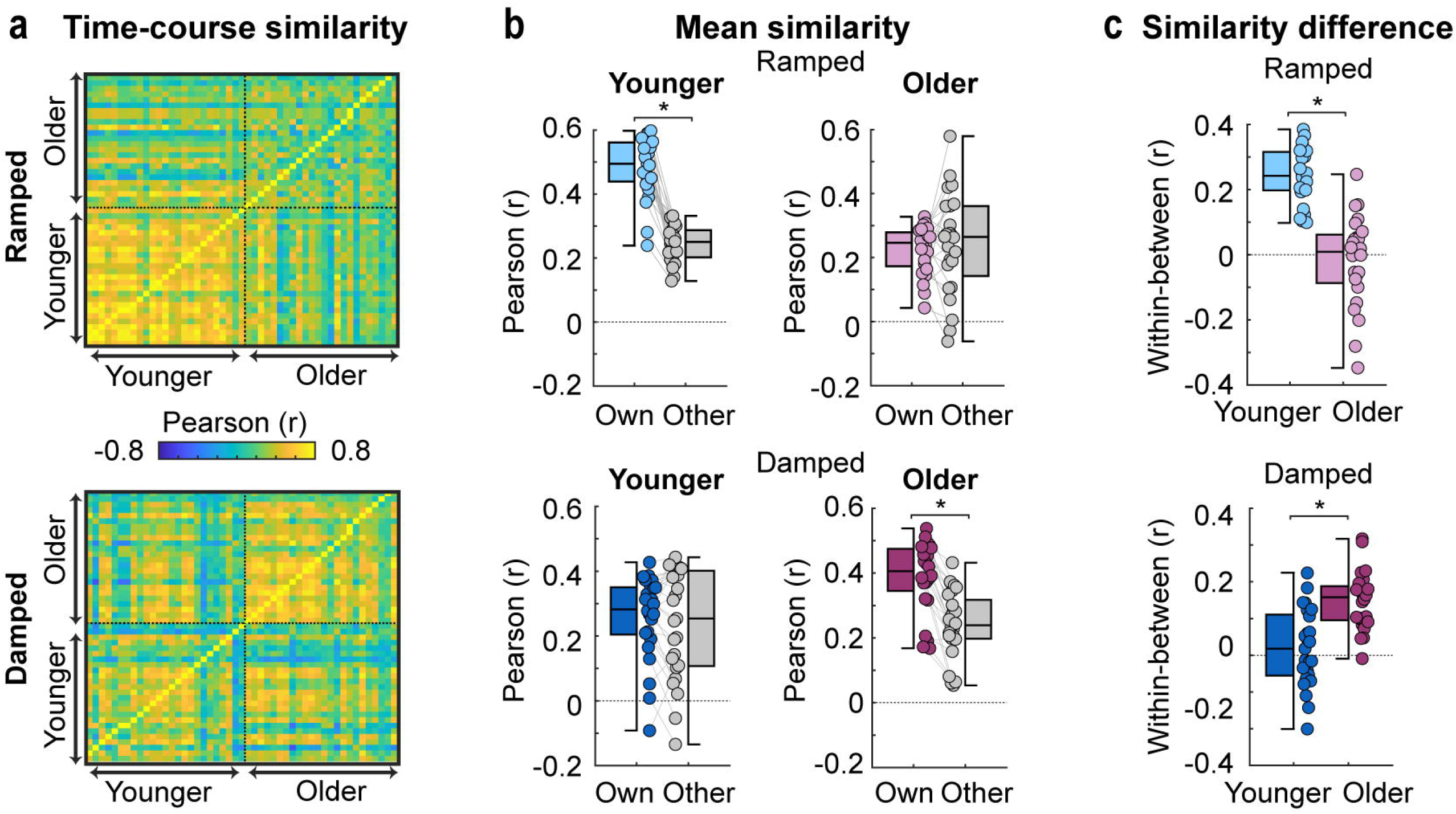
Results from the time-course correlation analysis. **(a)** Time-course similarity values (r) are plotted for ramped (top panel) and damped sounds (bottom panel). Each row contains *r* values representing an individual’s correlation between their own time course and the time course of every other individual in their own age group and other age group. **(b)** Mean similarity scores for ramped (top panels) and damped (bottom panels) are plotted for younger (left panels) and older subjects (right panels). Similarity scores are *r* values averaged based on group identity (own-group *r*, other-group *r*). **(c)** The difference in mean correlation (own-group *r* minus other group *r*) for each individual are plotted for ramped (top panels) and damped (bottom panels) and for younger and older subjects. Colored dots represent individual data points. Error bars reflect standard error. *p < 0.05.

Participants had larger own-group *r* compared to other-group *r* scores (effect of correlation type: *F*_1,47_ = 80.88, *p* = 8.8 × 10^-12^, η^2^_p_ = .63): participant time courses were more similar within an age group than across age groups. This difference between own-group and other group *r* was larger for younger compared to older subjects (correlation type × age group interaction: *F*_1,47_ = 9.05, *p* = 004, η^2^_p_ = .16). Similarity scores (*r*) did not differ as a function of envelope shape (*p* = .336), but a significant envelope shape × age group interaction (*F*_1,47_ = 30.59, *p* = 1.4 × 10^-6^, η^2^_p_ = .39), showed that younger subjects had larger similarity scores for *ramped* compared to damped (t_24_ = 4.24, *P_FDR_* = 6 × 10^-4^, r_e_ = .65), while older subjects had larger similarity scores for *damped* compared to ramped (t_24_ = −3.59, *P_FDR_* = .002, r_e_ = .65).

A significant envelope shape × correlation type × age group interaction (*F*_1,47_ = 90.1, *p* = 1.7 × 10^-12^, η^2^_p_ = .66) was analyzed with post-hoc t-tests. For ramped-envelope stimuli, younger participants had larger own-group *r* compared to other-group *r,* suggesting younger participants neural-response time courses to ramped envelopes were correlated more strongly among their peers than with time courses from older adults (t_24_ = 14.34, *P_FDR_* = 2.9 × 10^-13^, r_e_ = 0.95; Figure 5b, top left panel). Older participants showed no difference between own-group and other-group *r* for ramped-envelope stimuli *(p_FDR_* = .527; Figure 5b, top right panel). For damped-envelope stimuli, older adults had larger own-group *r* compared to other-group *r*, (t_23_ = 9.20, *P_FDR_* = 3.6 × 10^-9^, r_e_ = 0.89; Figure 5b, bottom right panel), while younger participants showed no difference between own- and other-group *r* for damped envelope shapes *(P_FDR_* = .369; Figure 5b, bottom left panel). Together, these analyses show that younger participants exhibit more synchronous neural responses when listening to *ramped envelopes* (slow onset and rapid offset) while older participants produce more synchronous neural responses when listening to *damped envelopes* (rapid onset and slow offset).

### Neural synchronization for different envelope shapes differs between younger and older adults

We quantified how envelope shape affects neural synchronization (ITPC) in older and younger adults (Figure 6a). For ITPC at the 4-Hz stimulation frequency, there was no effect of age group (*F*_1,47_ = 3.45, *p* = .01, η^2^_p_ =.07) nor envelope shape *(p* = .611), but the age group × envelope shape interaction was significant (*F*_1,47_ = 17.82, *p* = 1.10 × 10^-4^, η^2^_p_ = .28). Younger adults showed increased ITPC for sounds with ramped compared to damped envelope shapes (t_24_ = −3.27, *P_FDR_* = .006, r_e_ = 0.56), whereas older adults showed the reverse pattern (t_23_ = 2.69, *P_FDR_* = .013, r_e_ = 0.49; see Figure 6b). ITPC was also larger for sounds with high compared to low carrier-frequency bands for younger adults (t_24_ = −2.60, *p_FDR_* =.032, r_e_ = 0.47), but there was no difference for older adults *(p_FDR_* = .498; age group × carrier frequency interaction: *F*_1,47_ = 6.08, *p* = .017, η^2^_p_ = .12). There was a significant interaction between envelope shape and carrier frequency (*F*_1,47_ = 8.19, *p* = .006, η^2^_p_ = .15), but follow-up comparisons did not reveal any significant differences between ramped and damped low-carrier-frequency sounds *(p_FDR_* = .446) or ramped and damped high-carrier-frequency sounds *(p_FDR_* = .225). None of the other effects were significant (*F* < 2.67, *p* > .11, η^2^_p_ < .05).

ITPC for the fundamental/harmonic series (Figure 6b) did not differ between age groups *(p* = .678), but ITPC was larger for damped compared to ramped envelope shapes (effect of envelope shape: *F*_1,47_ = 13.55, *p* = .001, η^2^_p_ = .22). Critically, the envelope shape × age group interaction was significant (*F*_1,47_ = 7.01, *p* = .011, η^2^_p_ = .13; Figure 6b): neural synchronization was larger for damped compared to ramped envelope shapes for older adults (t_23_ = 5.35, *p_FDR_* = 3.9 × 10^-5^, r_e_ = 0.74), but did not differ between envelope shapes for younger participants *(p_FDR_* = .524). There was also an interaction between age group and carrier frequency (*F*_1,47_ = 6.02, *p* = .018, η^2^_p_ = .11). This was driven by reduced synchronization for sounds with a high compared to a low carrier frequency band in older participants (t_23_ = 3.17, *p_FDR_* = .046, r_e_ = 0.55), and a non-significant trend towards the opposite pattern in younger participants *(p_FDR_* = .269). None of the other effects were significant (*F* < 1.06, *p* > .31, η^2^_p_ < .02).

**Figure 6.**
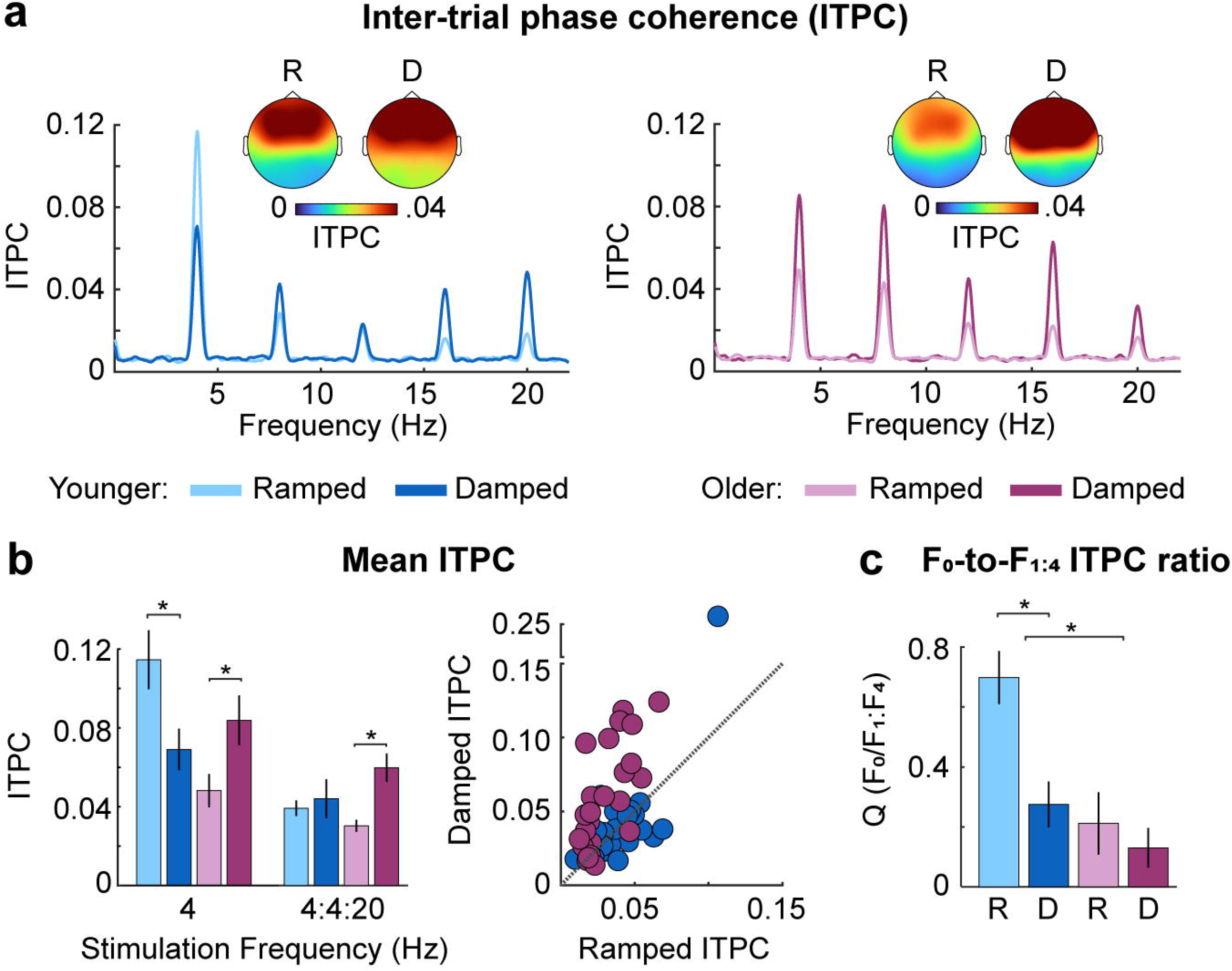
Results for neural synchronization. **(a)** Inter-trial phase coherence (ITPC), **(b)** mean ITPC at the stimulation frequency (4 Hz) and at the 4-Hz fundamental/harmonic series (4:4:20 Hz), and **(c)** the average ratio (Q) for synchronization to the stimulation frequency (F0) and the harmonics (Fl:F4) are plotted for each age group (younger, older) and envelope shape (ramped, damped). Topographies in (a) reflect mean ITPC at the 4-Hz stimulation frequency and are shown for each envelope shape (ramped, damped) and age group (younger, older). Error bars reflect standard error. *p < 0.05.

### Neural synchronization is less sinusoidal in older compared to younger adults

A growing body of evidence suggests that analyzing non-sinusoidal signal features can provide important information about underlying neural response properties (Cole and Voytek, 2017; Cole et al., 2017). To characterize the presence of non-sinusoidal signal features, we utilized the harmonic structure of the ITPC frequency spectrum to calculate *Q*, which indexes the degree to which the response is sinusoidal (spectral peak mainly at F0) versus non-sinusoidal (spectral peaks at the harmonics, F_1_ to F_4_). *Q* was smaller, indicating a less sinusoidal response, in older compared to younger participants (effect of age group: *F*_1,47_ = 9.67, *p* = .003, η^2^_p_ = .17) and was smaller (less sinusoidal response) for damped compared to ramped envelope shapes (effect of envelope shape: *F*_1,47_ = 15.54, *p* = 2.7 × 10^-4^, η^2^_p_ = .25). An interaction between age group and envelope shape (*F*_1,47_ = 7.13, *p* = .01, η^2^_p_ = .13) was due to younger participants having smaller *Q* for damped compared to ramped stimuli (t_24_ = 4.99, *P_FDR_* = 9 × 10^-5^, r_e_ = 0.71), whereas *Q* was small for both damped and ramped stimuli in older individuals, with no reliable difference (t_23_ = 0.85, *P_FDR_* = .406, r_e_ = 0.17). The synchronized response was therefore less sinusoidal for sounds with damped compared to ramped envelopes in younger adults, and non-sinusoidal for both envelope shapes in older adults (Figure 6c). There was an effect of carrier-frequency band: *Q* was smaller (less sinusoidal response) for sounds with a low compared to a high carrier-frequency band (*F*_1,47_ = 11, *p* = .002, η^2^_p_ = .19). Finally, there was an interaction between envelope shape and frequency band (*F*_1,47_ = 7.02, *p* = .011, η^2^_p_ = .13), which was a result of smaller *Q* for damped compared to ramped envelopes with a high carrier frequency band (t_48_ = 4.68, *P_FDR_* = 5 × 10^-5^, r_e_ = .56), but no difference between damped and ramped envelopes with a low frequency band (t_48_ = 1.87, *P_FDR_* = .067, r_e_ = .26). None of the other interactions were significant (*F* < 1.2, *p* > .29, η^2^_p_ < .02).

### Neural synchronization reflects sharper responses in older compared to younger adults

The fast Fourier transform decomposes a complex signal into sinusoids and may thus not be well suited to characterize non-sinusoidal features of neural responses. Analysis of specific aspects of the neural *signal shape,* such as sharpness, that is not well captured using the fast Fourier transform may provide important information about neural response properties and dysfunction (Sherman et al., 2016; Cole et al., 2017). In order to capture differences in neural signal shape we binned the amplitude data according to phase values of a 4-Hz sinusoid (Figure 7a), and then fit an exponential cosine function to the binned amplitude data (Figure 7b). We analyzed the estimated amplitude (*a*) and sharpness (exponent *e*) coefficients, with three-factor ANOVAs (envelope shape; carrier-frequency band; age group).

**Figure 7.**
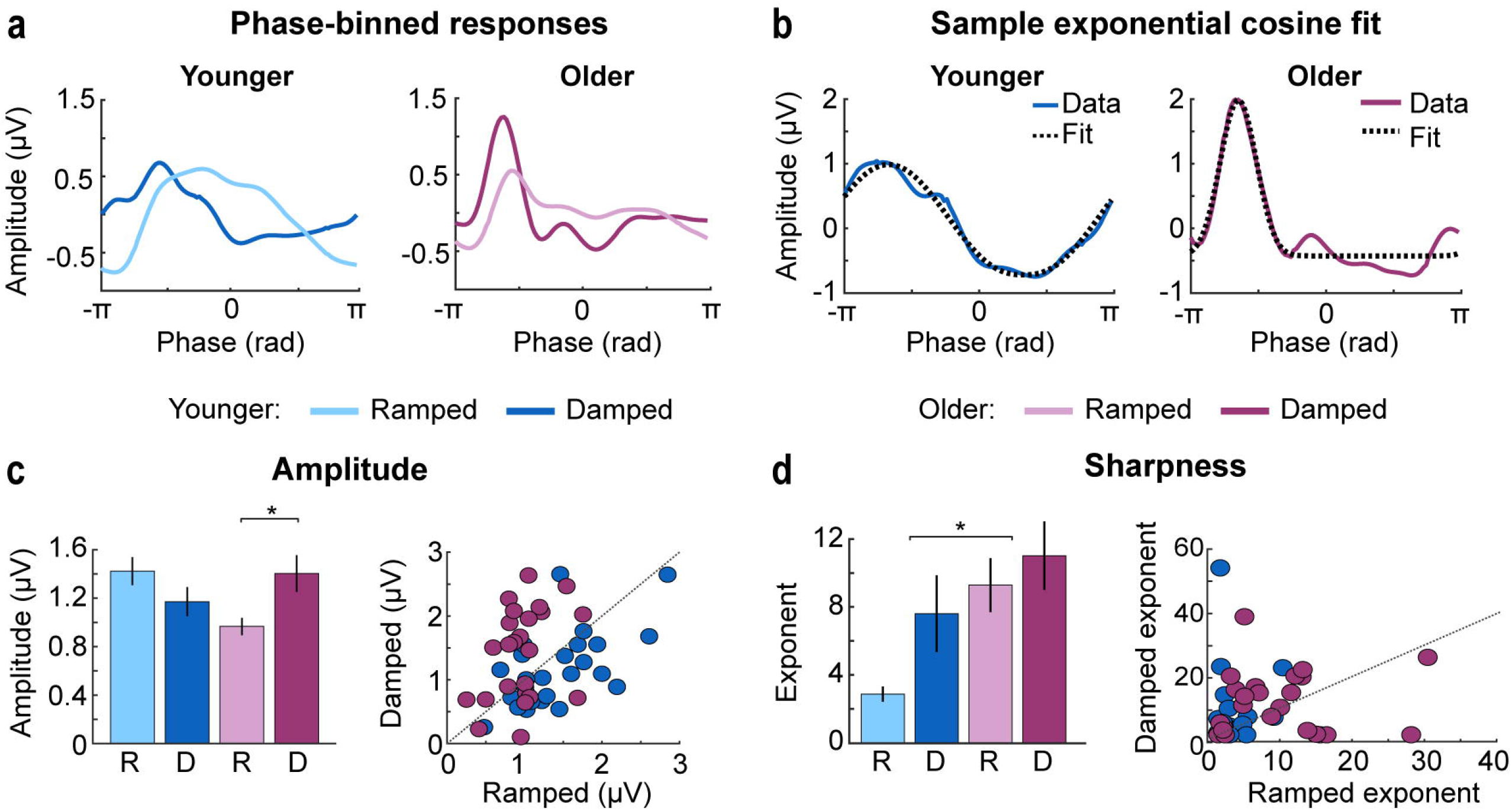
Results from the signal shape analysis. **(a)** Mean phase-binned amplitude values are plotted as a function of phase angle (radians) for younger (left panel) and older participants (right panel) and ramped and damped envelopes shapes. **(b)** A phase binned amplitude exemplar (Data) for younger (left panel) and older adults (right panel) is shown along with an example of an exponential cosine fit to these data (Fit). **(c)** Mean amplitude values are plotted as a function of age group (younger, older) and envelope shape (ramped, damped) in the left panel. Individual data points are shown on the right panel with a 45-degree reference line. **(d)** Mean exponent values reflecting the sharpness of the synchronized response are plotted as a function of age group (younger, older) and envelope shape (ramped, damped) in the left panel. Individual data points are shown on the right panel with a 45-degree reference line. Error bars reflect standard error. *p < 0.05.

The analysis of amplitude paralleled our ITPC findings (Figure 6b): there was no age group difference *(p* = .508), but amplitude was larger for damped compared to ramped envelope shapes (effect of envelope shape: *F*_1,47_ = 4.41, *p* = .041, η^2^_p_ = .09; Figure 7c). The interaction between age group and envelope shape was significant (*F*_1,47_ = 20.90, *p* = 3.5 × 10^-5^, η^2^_p_ = .31), revealing larger amplitudes for damped compared to ramped envelope shapes for older individuals (t_23_ = 4.48, *p_FDR_* = 3 × 10^-4^, r_e_ = 0.68), whereas a non-significant pattern in the opposite direction was observed for younger adults (t_24_ = −1.84, *p_FDR_* = .077, r_e_ = 0.35). There was also an interaction between age group and carrier-frequency band (*F*_1,47_ = 12.86, *p* = .001, η^2^_p_ = .22), which indicated that responses in younger participants were larger for high compared to low carrier-frequency sounds (t_24_ = 2.11, *p_FDR_* = .046, r_e_ = 0.40), whereas the reverse pattern was observed for older individuals (t_23_ = −3.11, *p_FDR_* = .01, r_e_ = 0.54). None of the other interactions were significant (*F* < 2.8, *p* > .1, η^2^_p_ < .06).

The exponent (indicating response sharpness) was larger for older compared to younger adults (effect of age group: *F*_1,47_ = 25.95, *p* = 6 × 10^-6^, η^2^_p_ = .36), and larger for damped compared to ramped envelopes (effect of envelope shape: *F*_1,47_ = 12.71, *p* = .001, η^2^_p_ = .21; Figure 7d). No remaining effects or interactions were significant (*F* < 2.8,*p* > .15, η^2^_p_ < .04). These results indicate that responses were sharper in older compared to younger adults, and for damped compared to ramped envelope shapes.

In sum, signal-shape analyses indicate that in older adults, responses were larger for envelopes with rapid onsets (and slow offsets) than for those with slow onsets (and rapid offsets), whereas younger adults showed the reverse pattern. Synchronized neural activity was less sinusoidal, and sharper, in older participants compared to younger participants.

### Relating neural sensitivity to different envelope shapes with measures of hearing loss

In order to quantify how neural sensitivity to different envelope shapes may be related to our measures of hearing loss, we calculated partial correlations (controlling for age) between PTA and the following neural measures: the difference in synchronization to damped versus ramped envelopes (4:4:20 Hz ITPC; averaged across carrier frequencies; *r*_46_ = −.09, *p_FDR_* = .556), the difference between damped and ramped *Q* scores (averaged across carrier frequencies; *r*_46_ = −.06, *p_FDR_* = .679), overall signal sharpness (averaged across envelope shapes and carrier frequencies; *r*_46_ = −0.02,*p_FDR_* = .918), and overall signal amplitude *a* (averaged across envelope shapes and carrier frequencies; *r*_44_ = −.23, *p_FDR_* = .117), but did not find any significant relationships. We also made a similar comparison between QuickSIN performance (0 SNR) and the same neural measures: the difference in synchronization to damped versus ramped envelopes (4:4:20 Hz synchronization: *r_46_* = −.09, *p_FDR_* = .67), the difference between damped and ramped *Q* scores (averaged across carrier frequencies; *r*_4_6 = −.13, *p_FDR_* = .607), overall signal sharpness (*r*_46_ = .02, *p_FDR_* = .916), and overall signal amplitude *a* (*r*_46_ = −.19,*p_FDR_* = .607, and, but no effects reached significance.

## Discussion

We examined neural sensitivity to sound envelopes with a ramped (gradual attack, sharp decay) or damped (sharp attack, gradual decay) envelope shape in younger and older adults. The three main findings are: (1) Auditory cortex of older adults is hyper-responsive to sound compared to younger adults despite similar subcortical responses in both groups; (2) Neural activity in older adults synchronizes more strongly with rapid-attack, slow-decay (damped) envelopes, whereas in younger adults it synchronizes more strongly with slow-attack, rapid-decay (ramped) envelopes; (3) Synchronized neural activity is less sinusoidal and sharper – appearing more burst-like – in older compared to younger adults. Our results demonstrate that older adults’ sensitivity to the amplitude envelope of sounds differs fundamentally from that of younger adults.

### Auditory cortex of older adults is hyper-responsive to sound

We observed larger cortical responses to sound onset in older compare to younger adults (Figure 4b), despite higher pure-tone thresholds (lower sensitivity) in older individuals (Figure 3a) and no difference in subcortical responses between age groups (Figure 3c). Our results are consistent with a growing literature showing hyper-responsiveness to sound in the cortex of older compared to younger rats (Hughes et al., 2010) and humans (Amenedo and Díaz, 1999; Snyder and Alain, 2005; Sörös et al., 2009; Lister et al., 2011; Alain et al., 2012; Herrmann et al., 2016, 2018; Herrmann and Butler, 2020), as well as in rats following noise exposure (Popelář et al., 1987; Syka et al., 1994; Manzoor et al., 2012) and adult humans with hearing loss compared to those without (Tremblay et al., 2003; Alain et al., 2014; Millman et al., 2017).

Hyper-responsiveness is thought to arise from damage to the auditory periphery, such that deprivation of inputs from peripheral structures to brain regions downstream leads to reduced neural inhibition and increased excitation throughout the auditory pathway (Caspary et al., 2008; Auerbach et al., 2014; Salvi et al., 2017). Consistent with the current results, studies comparing noise-exposed to control animals have repeatedly shown that hyper-responsiveness manifests most strongly in auditory cortex (Auerbach et al., 2014; Chambers et al., 2016; Asokan et al., 2018). In fact, hyper-responsiveness in auditory cortex has been taken as an index of a loss of inhibition (Ng and Recanzone, 2018). The enhanced onset responses in older compared to younger adults could also be a result of reduced response variability, as older adults have been shown to exhibit less variable neural response profiles compared to younger adults (Garrett et al., 2010, 2011). More consistent single-trial responses would result in a larger response magnitude in the average. Decreased response variability would likely be secondary to a loss of neural inhibition.

### Neural synchronization patterns differ between younger and older adults

By correlating each individual’s neural time course with that of other participants, we were able to derive a metric of neural response similarity for ramped and damped sounds across participants. This analysis is conceptually similar to calculating inter-subject correlation (Hasson et al., 2008; Cohen and Parra, 2016), and represents a measure of global synchrony with other participants. Time courses were more similar between individuals from the same age group than for individuals from different age groups, but this was only observed when participants listened to the envelope shape for which each age group showed heightened ITPC sensitivity. In younger individuals, neural-activity time courses were more synchronous with other younger participants than with older ones when listening to stimuli with a ramped envelope shape (Figure 5c). In contrast, older individuals exhibited neural-activity time courses for damped envelopes that were more synchronous with other older participants than with those of younger ones. Thus, older and younger subjects preferentially synchronize to specific envelopes shapes, and this effect is highly consistent across subjects.

Previous work has demonstrated larger synchronization strength at the stimulation frequency for older compared to younger adults (Goossens et al., 2016, 2018, 2019; Presacco et al., 2016b, 2016a). We did not observe an age-related increase in synchronization at the stimulation frequency, or when we additionally considered the energy at harmonic frequencies (4:4:20 Hz). Sound intensity has been shown to affect the magnitude of synchronized activity (Picton et al., 2003). Many studies control for audibility between normal-hearing and hearing-impaired participants by increasing the sound level for those with hearing impairment (Millman et al., 2017; Goossens et al., 2018, 2019). In some cases, a larger synchronization response in older compared to younger individuals is not observed if sounds are presented at the same level in both groups (Goossens et al., 2019). We used a sound level of ~75 dB SPL for both age groups, and observed that older adults were hyper-responsive to sound onset, but did not show increased synchronization with the AM stimulus, compared to younger listeners.

Compatible with previous findings in spiking activity of the inferior colliculus of rats (Herrmann et al., 2017), older adults showed increased synchronization strength for damped compared to ramped envelopes, while younger adults showed the opposite pattern (Figure 6b). The sizable response to damped compared to ramped envelopes across harmonics suggests older participants have hyper-sensitivity to sounds with sharp onsets. Further, we generally observed that the effects are consistent across both carrier frequency ranges, without consistent interactions between age group and carrier-frequency. The age-related changes in envelope sensitivity we observed appear to generalize across the range of frequencies tested here, which covers the critical frequency range used to discriminate speech sounds.

### The shape of synchronized neural activity differs fundamentally between age groups

Typically, neural synchronization with amplitude modulation focuses on the response at the stimulation frequency (sinusoidal component) (Purcell et al., 2004; Zhu et al., 2013; Bharadwaj et al., 2015; Dimitrijevic et al., 2016; Henry et al., 2017; Herrmann et al., 2017, 2019), although synchronization patterns commonly include non-sinusoidal response features, such as responses to the harmonics (Dallos, 1973; Lins et al., 1995; Cebulla et al., 2006; Zhu et al., 2013). Further, there is evidence that analyzing non-sinusoidal signal shape features – such as sharpness – can provide important physiological information about neural signaling and system dysfunction (Cole et al., 2017). We analyzed the extent to which neural responses consisted of primarily sinusoidal or non-sinusoidal response patterns by studying the ratio of responses at the fundamental to an average of the signal at the harmonics (*Q*). This showed that neural responses were overall less sinusoidal for damped envelopes compared to ramped, and for older compared to younger adults.

We also analyzed specific features of the neural signal shape and showed both ramped and damped envelopes elicited sharper neural responses in older compared to younger adults (Figure 7d), suggesting heightened sensitivity to both the sharp onset of damped envelopes and sharp offset of ramped envelopes. Sharp response features of neural activity indicates the response is likely driven by short synchronous bursts of activity: the same neural responses spread out in time would create a smoother more sinusoidal signal shape (Sherman et al., 2016; Cole and Voytek, 2017). These findings suggest that how the auditory system responds to amplitude envelopes in sounds is fundamentally changed in older individuals, such that responses are more synchronous, appearing burst-like.

### Relating envelope-shape sensitivity to hearing ability

Previous behavioral work has demonstrated that preserved envelope-shape cues are critical for speech intelligibility (Drullman et al., 1994; Shannon et al., 1995), and that poorer speech intelligibility in older individuals may be related to changes in envelope coding (Millman et al., 2017; Goossens et al., 2018). Despite observing fundamental changes in older adults’ neural sensitivity to amplitude envelopes, no relation with speech-in-noise ability was observed. One limitation of using the QuickSIN to assess speechin-noise here is that the babble noise masker has a nearly flat temporal profile, and may not reflect a reliance on temporal envelope cues as strongly as when using a speech-on-speech task, which has salient temporal fluctuations in the target and masker. Alternatively, observed changes in envelope sensitivity may be more related to other hearing difficulties that older adults experience, such as difficulty suppressing background sounds (Parmentier and Andrés, 2009; Mishra et al., 2014). Increased sensitivity to sharp stimulus features may increase the salience of background sounds with such features (e.g., stop consonants; Repp and Lin, 1989). Increased sensitivity to sharp stimulus features may also contribute to the discomfort some individuals experience when using a hearing aid with fast compression time constants (cf. Gatehouse et al., 2003), which distorts the temporal envelope, such that the attack is more rapid.

## Conclusions

We examined how different envelope shapes (ramped, damped) affect neural synchronization in younger and older adults. Older participants demonstrated neural hyper-responsiveness to sound onsets, despite showing no major differences in neural responses at peripheral and subcortical levels. Older participants also showed increased sensitivity to damped compared to ramped envelope shapes, whereas the opposite pattern was observed in younger adults. Furthermore, synchronized neural activity appeared less sinusoidal and more burst-like in older, compared to younger, individuals. Our findings underscore the importance of characterizing sinusoidal and non-sinusoidal features of synchronized neural responses to stimuli, and suggest that aging is accompanied by major changes in the way that brain activity synchronizes with amplitude modulations in sounds.

## Acknowledgements

This research was supported by the Canadian Institutes of Health Research (MOP133450 to I.S. Johnsrude). BH was supported by a BrainsCAN Tier I postdoctoral fellowship (Canada First Research Excellence Fund; CFREF) and the Canada Research Chair program.

